# Gut Microbiota in male patients with chronic traumatic complete spinal cord injury

**DOI:** 10.1101/417709

**Authors:** Chao Zhang, Wenhao Zhang, Jie Zhang, Yingli Jing, Mingliang Yang, Liangjie Du, Feng Gao, Huiming Gong, Liang Chen, Jun Li, Hongwei Liu, Chuan Qin, Yanmei Jia, Jiali Qiao, Bo Wei, Yan Yu, Hongjun Zhou, Zhizhong Liu, Degang Yang, Jianjun Li

**Affiliations:** School of Rehabilitation Medicine, Capital Medical University, Beijing 100068, China; Department of Spinal and Neural Function Reconstruction, China Rehabilitation Research Center, Beijing 100068, China; Center of Neural Injury and Repair, Beijing Institute for Brain Disorders, Beijing 100068, China; China Rehabilitation Science Institute, Beijing 100068, China; Beijing Key Laboratory of Neural Injury and Rehabilitation, Beijing 100068, China; Institute of Rehabilitation medicine, China Rehabilitation Research Center, Beijing 100068, China; Department of Spinal Cord Injury Rehabilitation, China Rehabilitation Research Center, Beijing 100068, China; Laboratory medicine, China Rehabilitation Research Center, Beijing 100068, China

**Author notes:** **Corresponding authors:** 1 Jianjun Li and 2 Degang Yang. School of Rehabilitation Medicine, Capital Medical University, No. 10 Jiaomen North Road, Fengtai District, 100068 Beijing, China;. **Author Disclosure Statement** No competing interests exist.

**Keywords:** gut microbiota dysbiosis, chronic traumatic complete SCI, neurogenic bowel management, NBD symptoms, serum biomarkers

## Abstract

This study examined the diversity and structure of gut microbiota in healthy adults and chronic traumatic complete spinal cord injury (SCI) patients, documented neurogenic bowel management of SCI patients. The V3-V4 region of 16S rRNA gene from DNA of 91 fecal samples of 48 healthy and 43 diseased subjects was amplified and sequenced. There was difference in gut microbiota between healthy adult males and females. Neurogenic bowel dysfunction (NBD) was common in patients with chronic traumatic complete SCI, patients with quadriplegia have longer time to defecate than paraplegic patients, with higher NBD scores and heavier neurogenic bowel symptoms. Gut microbiota dysbiosis existed in SCI patients. The abundance of Veillonellaceae and Prevotellaceae increased while Bacteroidaceae and Bacteroides decreased in SCI group. The abundance of Bacteroidaceae, Bacteroides in quadriplegia group and Acidaminococcaceae, Blautia in paraplegia group were significant high than the health male group. Serum biomarkers GLU, HDL, CR and NBD symptoms defecation time, COURSE had significant correlation with microbial community structure. This study presents a comprehensive landscape of gut microbiota in adult male patients with chronic traumatic complete SCI and documents their neurogenic bowel management. The gut microbiota dysbiosis of SCI patients was correlation with serum biomarkers and NBD symptoms.

**IMPORTANCE:** Neurogenic bowel dysfunction is a major physical and psychological problem in patients with spinal cord injury, which can seriously affect the quality life of them. Gut dysbiosis are highly likely to occur in spinal cord injury patients There are few studies on intestinal microecology after spinal cord injury, and the clinical studies are fewer. It is importance to document their neurogenic bowel management and present a landscape of gut microbiota in them. We found the gut microbiota dysbiosis of spinal cord injury patients was correlation with serum biomarkers and neurogenic bowel dysfunction symptoms. These results may have implications in the next study about metagenomics and precision treatment of neurogenic bowel dysfunction in spinal cord injury patients.

## Introduction

After complete spinal cord injury, the loss of descending control over sympathetic preganglionic neurons causes autonomic reflex circuitry to become dysfunctional creating pathology including autonomic dysreflexia and SCI–immune depression syndrome (1,2,3,4,5), it causes an autonomic imbalance in the gastrointestinal tract, which leads to deficits in colonic motility, mucosal secretions, and vascular tone (6,7). The early survival rate of such patients has been significantly improved, but the quality of life of such patients is still not satisfactory. Among them, neurogenic bowel dysfunction (NBD) is a major physical and psychological problem in patients with SCI, which can seriously affect the quality life of patients. The two main manifestations of NBD are constipation and fecal incontinence, with the prevalence of constipation in these patients reported to be 40–58%, and fecal incontinence from 2 to 61% (8,9,10,11). Because of these problems, patients with chronic SCI tend to spend more time in the toilet while evacuating their bowels, use suppositories, laxatives and supplemental dietary fiber more frequently to improve bowel evacuation and require manual removal of feces much more frequently when compared with their matched control population (12,13,14,15). One of the aims of our study was to document neurogenic bowel management of chronic traumatic completed SCI male patients in our center.

Human intestinal tract is colonized by thousands of different genera of bacterial species whose number and genetic content exceed that of the host by a factor of ten and 150-fold, respectively (16). That is critical for normal digestion, nutrient absorption, and the development, metabolism, and function of cells throughout the body (17,18). Recent studies have shown that an imbalance of the normal gut microbiota (dysbiosis) is associated with inflammatory bowel diseases (19), irritable bowel syndrome and some other diseases (20,21).

Elin O et al reported that sex hormones affected the gut microbiota composition in male and female mice in a controlled environment; Francesca Borgo et al reported that body mass index and gender affect microbial flora in different parts of the gut (22,23). One of the aims of this study was to explore whether there is a difference of gut microbiota in healthy adult males and females.

Common causes of gut dysbiosis include antibiotic use, prolonged stress, and gastrointestinal dysfunction (17,24,25). Because most patients with acute complete SCI have changed the intestinal transit time and destructed the intestinal mucosal function barrier after injury, the displacement of the intestinal flora making the intestines to be the largest “endotoxin pool” in the human body. The use of antibiotics must affect healthy intestinal micro-ecological systems (26,27,28,29). Therefore, gut dysbiosis are highly likely to occur in SCI.

There are few studies on intestinal microecology after SCI in clinical studies. Kigerl KA et al have shown that traumatic SCI can cause intestinal disorders, and that dysbiosis can impair functional recovery through stool samples from traumatic SCI mice (30). Bilgi Gungor et al reported a clinical study of 30 patients with SCI, showed that the number of butyrate communities in patients with SCI was significantly lower than that in the normal population (29). More researches are needed to determine whether intestinal dysbiosis after SCI changes in a range of clinically relevant variables (31).

The study of this article is to explore the difference of healthy adult males and females in gut microbiota, document neurogenic bowel management of chronic traumatic complete SCI male patients in our center; To investigate the comparative analysis of intestinal gut microbiota in chronic traumatic complete SCI male patients and healthy males. Exploring the association between intestinal microbiota with serum biomarkers and neurogenic bowel symptoms.

## Results

### Baseline characteristics of the samples in the health male and female groups

The mean age for 23 healthy adult males and 25 females was 40±9 years and 37±8 years (18–60 years), there was no statistically significant differences (one-way ANOVA, p=0.255). The BMI in males was significantly high (24.8±2.777) than females (22.8±2.763), (one-way analysis of variance, p = 0.015).

### Diversity and taxonomic analysis in the health male and female groups

16S rRNA gene sequences were generated using Illumina’s MiSeq platform. Briefly, a total of 1010832 sequences were obtained. Reads were clustered in OTUs at 97% of identity. The rarefaction curves showed clear asymptotes and the Good’s coverage for the observed OTUs was 99.46%, which together indicate a near-complete sampling of community. No significant difference in either OTU abundance or OTU diversity index was observed between the male and female populations (Fig. 1A and Supplementary File 1). The discrete case of sample points distribution in PLS-DA on genus level showed differences in the composition of the gut microbiota between the two groups (Fig. 1B).

**Figure 1.**
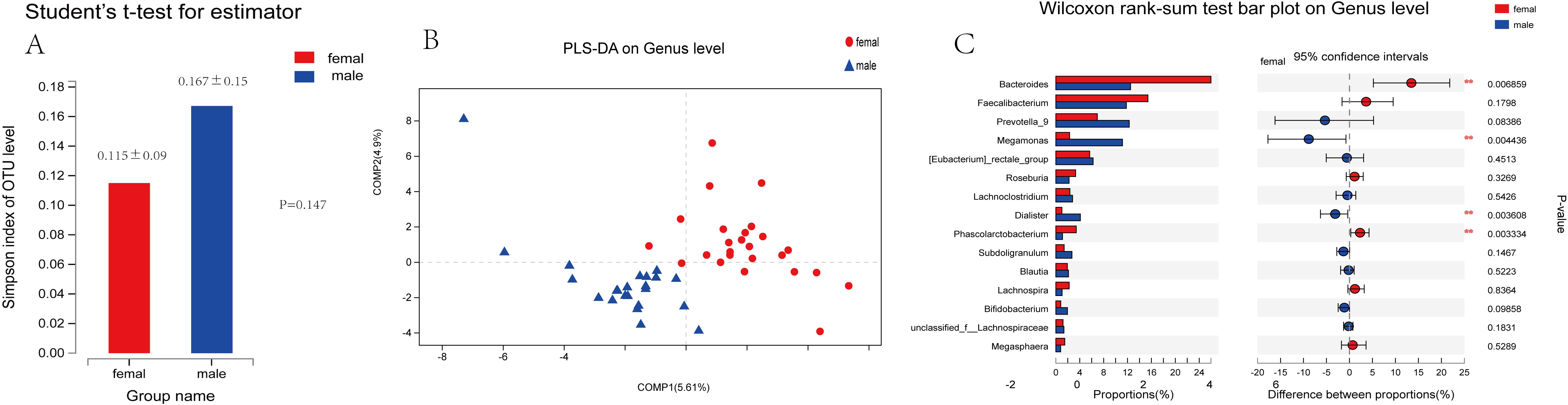
Diversity and taxonomic analysis in the health male and female groups A: No significant difference in OTU abundance (Simpson index) was observed between the male and female populations (p=0.147). B: The discrete case of sample points distribution in PLS-DA on Genus level showed differences in the composition of the gut flora between male and female groups. C: Genus-level operational taxonomic units different between healthy male and female groups. STAMP software was used to calculate the genus proportions in the two groups. There were 4 of top 15 genus showed a significant difference (P<0.05)among two groups (Welch’s t-test).

STAMP analysis indicates there were 15 OTUs showed a significant difference (P<0.05)among two groups (Welch’s t-test). There were 4 of top 15 genus showed a significant difference(P<0.05)among two groups (Welch’s t-test). The abundance of Megamonas and Dialister in male group were significant high than the female group (P< 0.01, P< 0.01, Mann-Whitney U test); the abundance of Bacteroides and Phascolarctobacterium in female group were significant high than the male group (P< 0.01, P< 0.01, Mann-Whitney U test) (Fig.1C). We can find that there was a difference in fecal flora between healthy adult males and females.

### Characteristics and neurogenic bowel management of male patients with chronic traumatic complete SCI

In all, 43 patients with chronic SCI fulfilling the enrollment criteria were interviewed and completed the survey form (Table 1). The causes of injury were traffic accidents (37.2%), bruised by heavy object (20.9%), fall from height (20.9%), in that order. The mean score of NBD was 10.02±5.11. The mean defecation time was 35.33±16.766 minutes. Most patients (60.5%) took bowel care not daily but more than twice every week, the others (39.5%) frequency of bowel care were once daily. Main techniques for fecal evacuation was suppository (88.4%), manual evacuation (23.3%), digital stimulation (16.3%), spontaneous (4.7%), in that order. Supplementary interventions for fecal evacuation were abdominal massage (58.1%), digital anus-rectal stimulation (48.8%), digital evacuation (9.4%), taking cathartic drug (9.4%).

**Table 1.** Neurogenic Bowel Management table in male Patients with chronic traumatic completed SCI

|  | SCI-Male n (%) | SCI-cervical n (%) | SCI-thoracic and lumbar n (%) | P |
| --- | --- | --- | --- | --- |
| Course | 62.5±53.98 | 69.4±52.72 | 56.5±54.36 | 0.449 |
| NBD Scores | 10.02±5.11 | 11.17±5.16 | 8.7±4.72 | 0.119 |
| Defecation time | 35.33±16.766 | 41.789±19.29 | 30±13.94 | 0.026 |
| Pathogenesis | Traffic accident<br>16 (37.2%)<br>Bruised by heavy object<br>9 (20.9%)<br>Falling down<br>9(20.9%)<br>Other causes<br>9(20.9%) | Traffic accident<br>10 (50%)<br>Bruised by heavy object<br>4(20%)<br>Falling down<br>3(15%)<br>Other causes<br>3(15%) | Traffic accident<br>6 (26.1%)<br>Bruised by heavy object<br>5 (21.7%)<br>Falling down<br>6 (26.1%)<br>Other causes<br>6 (26.1%) |  |
| Frequency of bowel care | Once a day:<br>17 (39.5%)<br>Not daily but more than twice every week:<br>26 (60.5%) | Once daily:<br>5 (25%)<br>Not daily but more than twice every week<br>15 (75%) | Once daily:<br>12 (52.2%)<br>Not daily but more than twice every week<br>11 (47.8%) |  |
| Main techniques for faecal evacuation | Suppository<br>38 (88.4%)<br>Digital stimulation<br>7 (16.3%)<br>Manual evacuation<br>10 (23.3%)<br>Spontaneous<br>2 (4.7%) | Suppository<br>20 (100%)<br>Digital stimulation<br>6 (30%)<br>Manual evacuation<br>8 (40%) | Suppository<br>18 (78.3%)<br>Digital stimulation<br>1 (2.3%)<br>Manual evacuation<br>2 (4.6%)<br>Spontaneous<br>2 (4.6%) |  |
| Supplementary interventions | Abdominal massage<br>25 (58.1%)<br>Digital anus-rectal stimulation<br>21 (48.8%)<br>Digital evacuation<br>4 (9.3%)<br>taking cathartic drug<br>4 (9.3%) | Abdominal massage<br>10 (50%)<br>Digital anus-rectal stimulation<br>7 (35%)<br>Digital evacuation<br>2 (10%)<br>taking cathartic drug<br>1 (5%) | Abdominal massage<br>15 (65.2%)<br>Digital anus-rectal stimulation<br>14 (60.9%)<br>Digital evacuation<br>2 (8.7%)<br>taking cathartic drug<br>3 (13%) |  |
| Timing of bowel care | Morning 4 (9.3%)<br>Afternoon 27 (62.8%)<br>Evening 9 (20.9%)<br>Inconsistent 3 (7%) | Morning 1 (5%)<br>Afternoon 14 (70%)<br>Evening 5 (25%) | Morning 3 (13%)<br>Afternoon 13 (56.5%)<br>Evening 4 (17.4%)<br>Inconsistent 3 (13%) |  |
| Location during | Bed | Bed | Bed |  |
| evacuation | 19 (44.2%)<br>toilet seat<br>8 (18.6%)<br>Potty chair<br>16 (37.2%) | 12 (60%)<br>toilet seat<br>2 (10%)<br>Potty chair<br>6 (30%) | 7 (30.4%)<br>toilet seat<br>6 (26.1%)<br>Potty chair<br>10 (43.5%) |  |
| Degree of assistance needed | Need all help<br>23 (53.5%)<br>Need partial help<br>10 (23.3%)<br>Independent completion<br>9 (20.9%)<br>Need special help<br>1 (2.3%) | Need all help<br>19 (95%)<br>Need special help<br>1 (5%) | Need all help<br>4 (17.4%)<br>Need partial help<br>10 (43.5%)<br>Independent completion<br>9 (39.1%) |  |
| Abdominal discomfort | 27 (62.8%) | 12 (60%) | 15 (65.2%) |  |
| Constipation | 29 (67.4%) | 14 (70%) | 15 (65.2%) |  |
| Bloating symptom | 32 (74.4%) | 16 (80%) | 16 (69.6%) |  |
| Flatus incontinence | 38 (88.4%) | 18 (90%) | 20 (87%) |  |
| Lifestyle alteration due to NBD | Major impact<br>25 (58.1%)<br>Some impact<br>15 (43.9%)<br>Little impact<br>3 (7%) | Major impact<br>12 (60%)<br>Some impact<br>6 (30%)<br>Little impact<br>2 (10%) | Major impact<br>13 (56.5%)<br>Some impact<br>9 (39.1%)<br>Little impact<br>1 (4.3%) |  |
| Top 3 complication desired to be solved | Neurogenic bowel dysfunction<br>42 (97.7%)<br>neurogenic bladder<br>36 (83.7%)<br>Sexual Dysfunction<br>19 (44.2%)<br>spasm<br>14 (32.6%)<br>neuralgia<br>8 (18.6%) | Neurogenic bowel dysfunction<br>19 (95%)<br>neurogenic bladder<br>16 (80%)<br>Sexual Dysfunction<br>8 (40%)<br>Spasm<br>7 (35%)<br>neuralgia<br>3 (15%) | Neurogenic bowel dysfunction<br>23 (100%)<br>neurogenic bladder<br>20 (87%)<br>Sexual Dysfunction<br>11 (47.8%)<br>Spasm<br>7 (30.4%)<br>neuralgia<br>5 (21.7%) |  |

**Table2.** Demographics and serum biomarkers between male healthy and patients with chronic traumatic completed SCI

|  | Health male | SCI-Male | P |
| --- | --- | --- | --- |
| N | 23 | 43 |  |
| AGE | 40±9.03 | 39.9±10.57 | 0.998 |
| BMI | 24.8±2.677 | 23.11±2.876 | 0.022 |
| ALT | 26.791±16.367 | 26.2±19.303 | 0.903 |
| AST | 23.848±17.097 | 21±9.8 | 0.429 |
| GLU | 4.343±0.528 | 5.266±1.964 | 0.033 |
| TG | 1.436±1.319 | 1.928±1.207 | 0.137 |
| TCHO | 3.695±0.794 | 4.217±1.005 | 0.038 |
| HDL | 0.9152±0.2091 | 0.917±0.163 | 0.974 |
| LDL | 2.177±0.596 | 2.617±0.701 | 0.005 |
| UREA | 4.416±1.224 | 4.403±1.14 | 0.966 |
| CR | 64.3±12.701 | 60.7±11.8 | 0.265 |
| UA | 309±69.81 | 378.1±64.93 | 0.001 |

More than a half patients bowel care time was in afternoon (62.8%), the other patients’ bowel care time was in evening (20.9%), and in morning (9.4%). The location of bowel care was bed (44.2%), potty chair (37.2%) and toilet seat (18.8%). 53.5% patients need all help during the defecation time, 25.6% patients need partial help, 20.9% patients can defecation independent. 62.8% patients had an abdominal discomfort symptom, 67.4% patients had a constipation symptom, 74.4% patients had a bloating symptom, 88.4% patients had flatus incontinence. The most common top 3 complications that patients wanted to solved were neurogenic bowel dysfunction(100%), neurogenic bladder(83.7%), sexual dysfunction(44.2%).

The quadriplegia SCI patients had a significant high BMI (23.586±3.35) than paraplegia SCI patients (22.697±2.31) (P<0.001). There were statistical differences between the two groups in HDL UREA and CRP (P<0.001) (Table3). Compared with paraplegia SCI patients, the quadriplegia SCI patients had longer defecation time, higher NBD score, lower defecation frequency, need more supplementary interventions to complete the bowel care.

Most of the defecation locations in the quadriplegia SCI patients were in the bed. Almost all the quadriplegia SCI patients require total help to complete the bowel care, but most paraplegia SCI patients could finish the bowel care independently or only need partially help. Most of SCI patients have abdominal discomfort such as constipation, bloating, and flatus incontinence. More than half of the patients in the two groups have a serious impact on lifestyle, and the most common complication they want to resolve is NBD.

### Composition of the gut microbiome of health male group and male chronic traumatic complete SCI groups

To exclude the effect of gender on gut microbiota results, we selected 23 male healthy individuals and 43 male patients with spinal cord injury to perform a comparative analysis. Demographics and serum biomarkers between male healthy and patients with chronic traumatic completed SCI were showed in Table2.

Briefly, a total of 2247802 sequences were obtained. Reads were clustered in OTUs at 97% of identity. The rarefaction curves showed clear asymptotes and the Good’s coverage for the observed OTUs was 99.88%, which together indicate a near-complete sampling of community. 798 OTUs are recognized in total. No significant difference in OTU abundance (ace, chao1 index) was observed between health male and SCI populations. In genus level, OTU diversity index Simpson showed a significant difference between two groups (P=0.03635)(Fig.2A). This indicates a decrease in intestinal flora diversity in patients with SCI.

**Figure 2.**
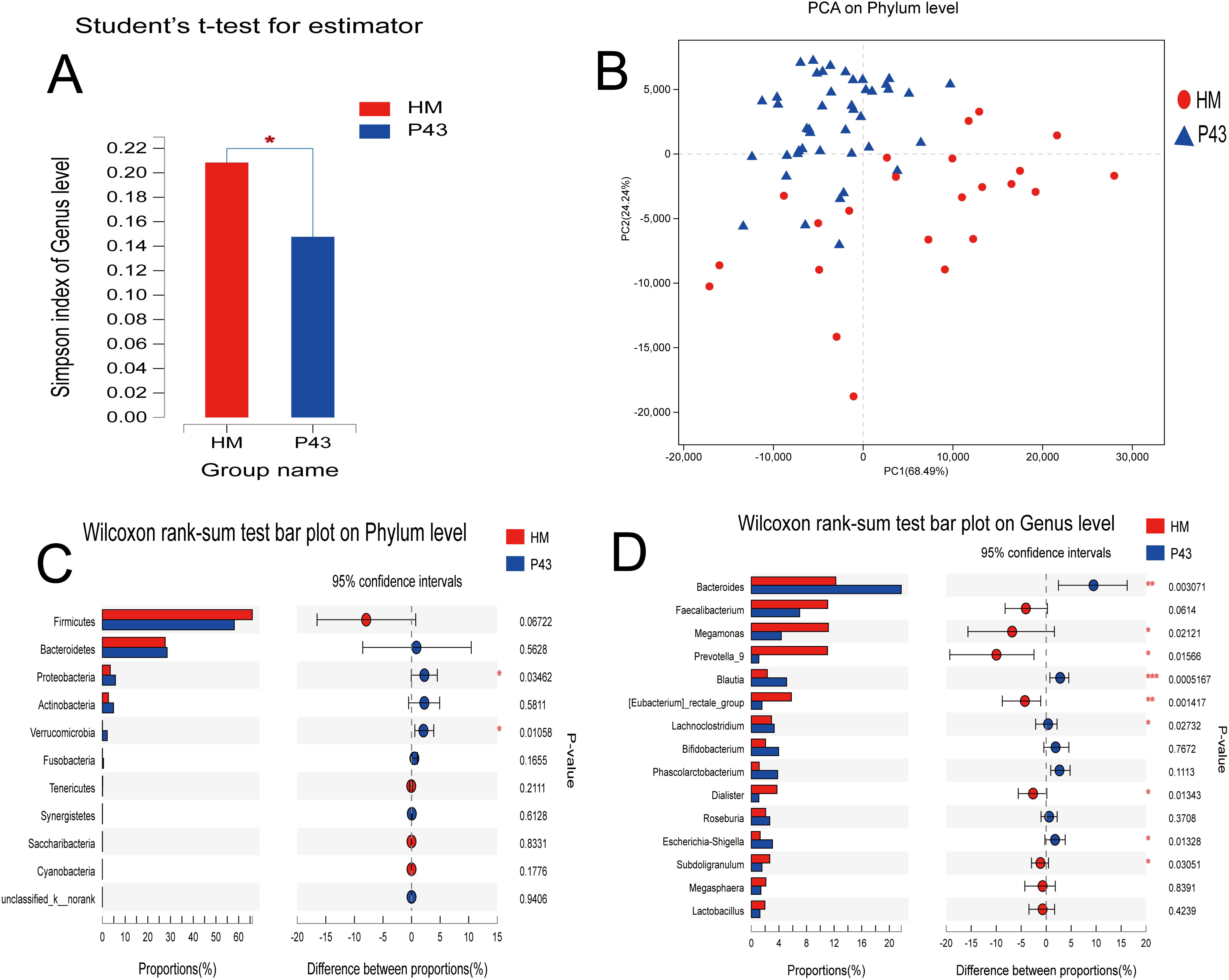
Diversity and taxonomic analysis in the health male and SCI groups A: In genus level, Simpson index showed a significant difference between healthy male and SCI groups (P=0.03635). B. Plot of principal coordinate analysis (PCA) on Phylum level of the fecal microbiota based on the unweighted UniFrac metric in healthy male and SCI groups. STAMP analysis on Phylum and Genus level showed differences between healthy male and SCI groups. There were 2 of top 15 phylum (C) and 9 of top 15 genus (D) showed a significant difference(P<0.05)among two groups (Welch’s t-test).

The PCA on phylum level and the NIMDS on OTU and genus level of beta-diversity analysis showed there were significant differences in bacterial community composition between two groups. ANOSIM/Adonis revealed significant differences in the structure of gut microbiota among the two groups (p<0.05) (Supplementary File 2). PLS-DA revealed that there were significant differences in bacterial community composition between two groups on OTU, phylum and genus level (p < 0.05) (Fig.2B).

STAMP analysis indicates there were 9 of top 15 genus showed a significant difference(P < 0.05)among two groups (Welch’s t-test). The abundance of Megamonas, Prevotella_9, (Eubacterium)_rectale_group, Dialister, Subdoligranulum in male group were significant high than the SCI group (p<0.05, Mann-Whitney U test); the abundance of Bacteroides Blautia Lachnoclostridium Escherichia-Shigella in SCI group were significant high than the male group (P< 0.05, Mann-Whitney U test)(Fig.2C, D). By LEfSe analysis (LDA threshold of 2), it was found that Veillonellaceae and Prevotellaceae were significantly enriched in SCI group compared with Bacteroidaceae and Bacteroides enriched in healthy male group.

According to NBD constipation symptom, we divided the SCI patients into constipation group and without constipation group. STAMP analysis showed a significant difference (P<0.05) among two groups (Welch’s t-test) in Bifidobacterium on Genus level (Fig.3A). We also divided the SCI patients into bloating group and without bloating group according to bloating symptom, STAMP analysis showed Megamonas had a significant high (P<0.05) in bloating group and Alistipes had a significant high (P<0.05) in without bloating group on genus level (Fig.3B).

**Figure 3.**
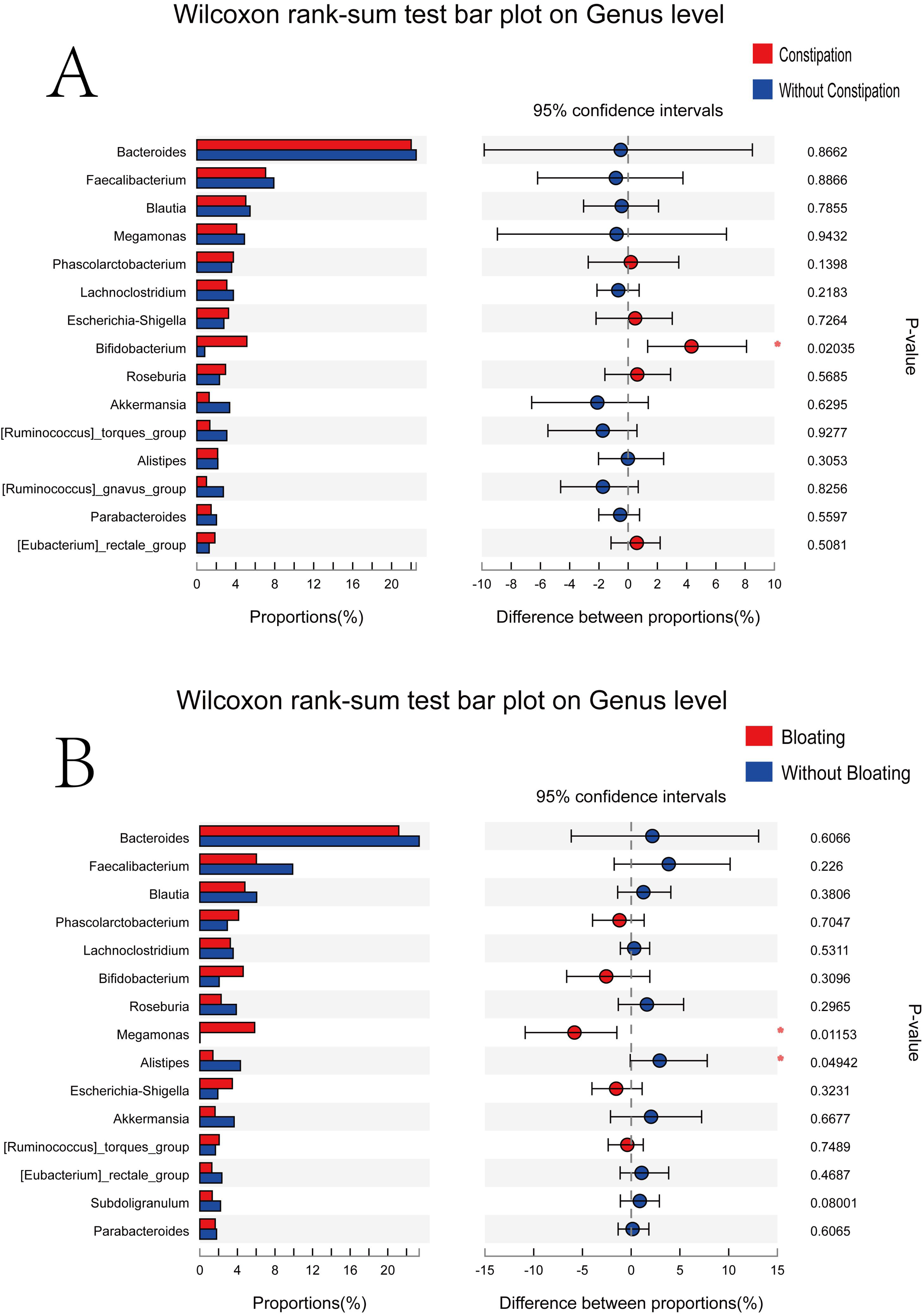
STAMP analysis on NBD symptoms A. STAMP analysis showed a significant difference (P < 0.05)among two groups (Welch’s t-test) in Bifidobacterium on Genus level. B.STAMP analysis showed Megamonas had a significant high(P < 0.05)in bloating group and Alistipes had a significant high(P < 0.05)in without bloating group on Genus level.

The selected environmental factors: BMI, AGE, ALT, AST, GLU, TG, TCHO, HDL, LDL, UREA, CR, UA for RDA analysis. One-way ANOVA showed statistically significant differences in BMI, GLU, TCHO, LDL and UA between the two groups (P < 0.05). RDA/CCA showed that GLU (p=0.017, r^2^=0.1315) HDL (p=0.028, r^2^=0.1121) CR (p=0.017, r^2^=0.1349) significantly affected bacterial composition in phylum level; In top 20 genus, BMI (p=0.04, r^2^=0.0971), GLU (p=0.044, r^2^=0.108)and HDL (p=0.001, r^2^=0.3044) significantly affected bacterial composition. We can found that Serum biomarkers GLU HDL and CR had significant correlation with microbial community structure (p<0.05).

Correlation heatmap analysis of different environmental factors on the community composition of two groups showed that Proteobacteria was positively correlated with UA (Pearson r=0.26, p=0.035); Cyanobacteria were positively correlated with AST (Pearson r=0.355, p=0.003); Fusobacteria were negatively correlated with AGE (Pearson r=-0.342, p=0.005) in phylum level(Fig.4A). In top 20 genus, Bacteroides was negative correlated with HDL (Pearson r=-0.418, p<0.001); Megamonas was negatively correlated with GLU (Pearson r=-0.513, p<0.001); Blautia was positively correlated with UA (Pearson r=0.274, p=0.026); Dialister was negatively correlated with UA, LDL, TG and TCHO (Pearson r=-0.32, P=0.009; r=-0.289 P=0.019; r=-0.258, P=0.037; r=-0.303, P=0.013 respectively.) (Fig.4B and Supplementary File 3-4).

**Figure 4.**
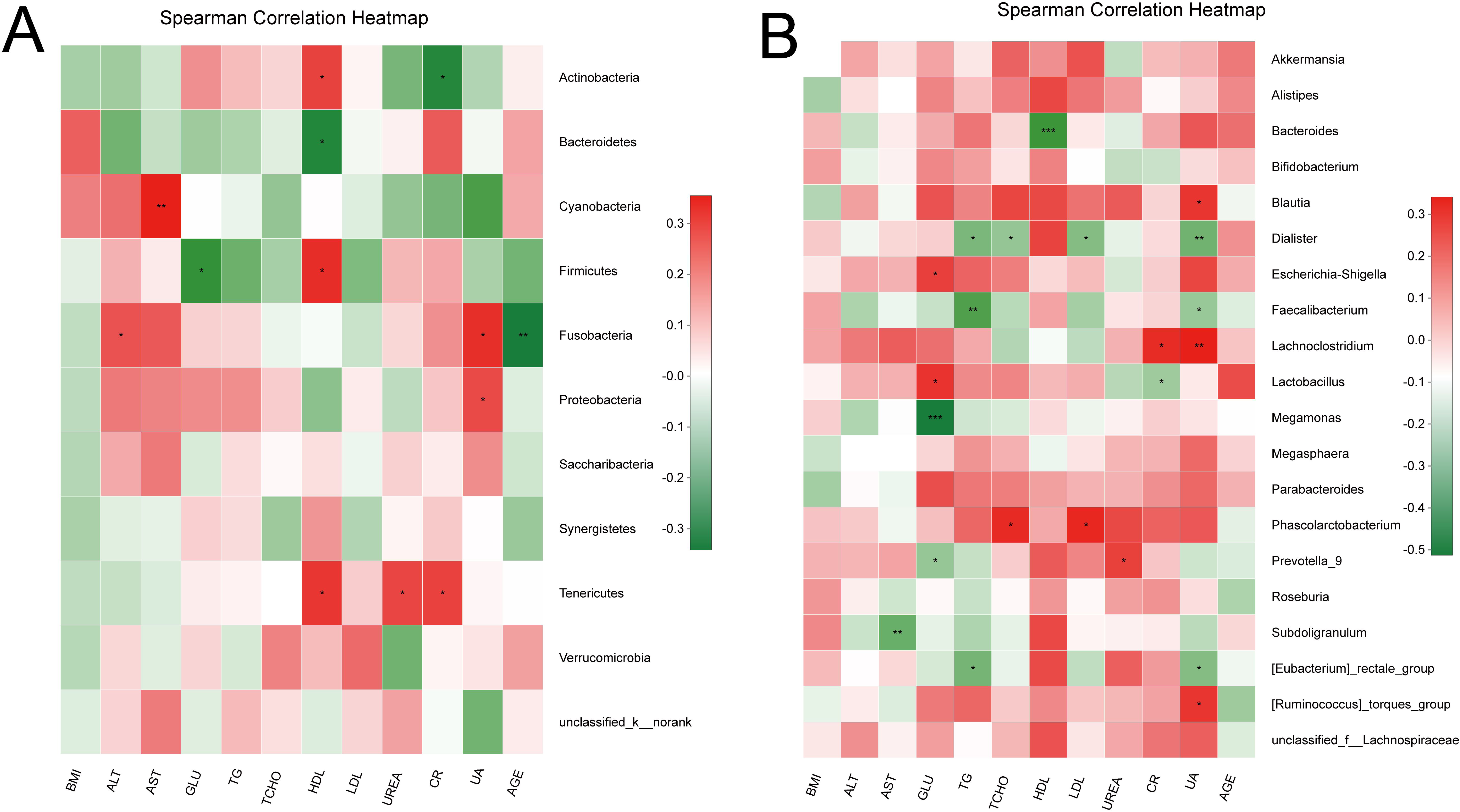
Correlation heatmap analysis of different environmental factors on the community composition of the healthy male and SCI groups in phylum level (A) and genus level(B).

### Comparison of the gut microbiome in quadriplegia and paraplegic groups

We divided the 43 SCI patients into 20 quadriplegia group and 23 paraplegic group, the characteristics and neurogenic bowel management were showed in Table 1 and 3. We found that the defecation time of quadriplegia patients (41.789±19.29minutes) was significant high than paraplegic patients (30±13.94minutes) (P=0.026).

**Table3.**
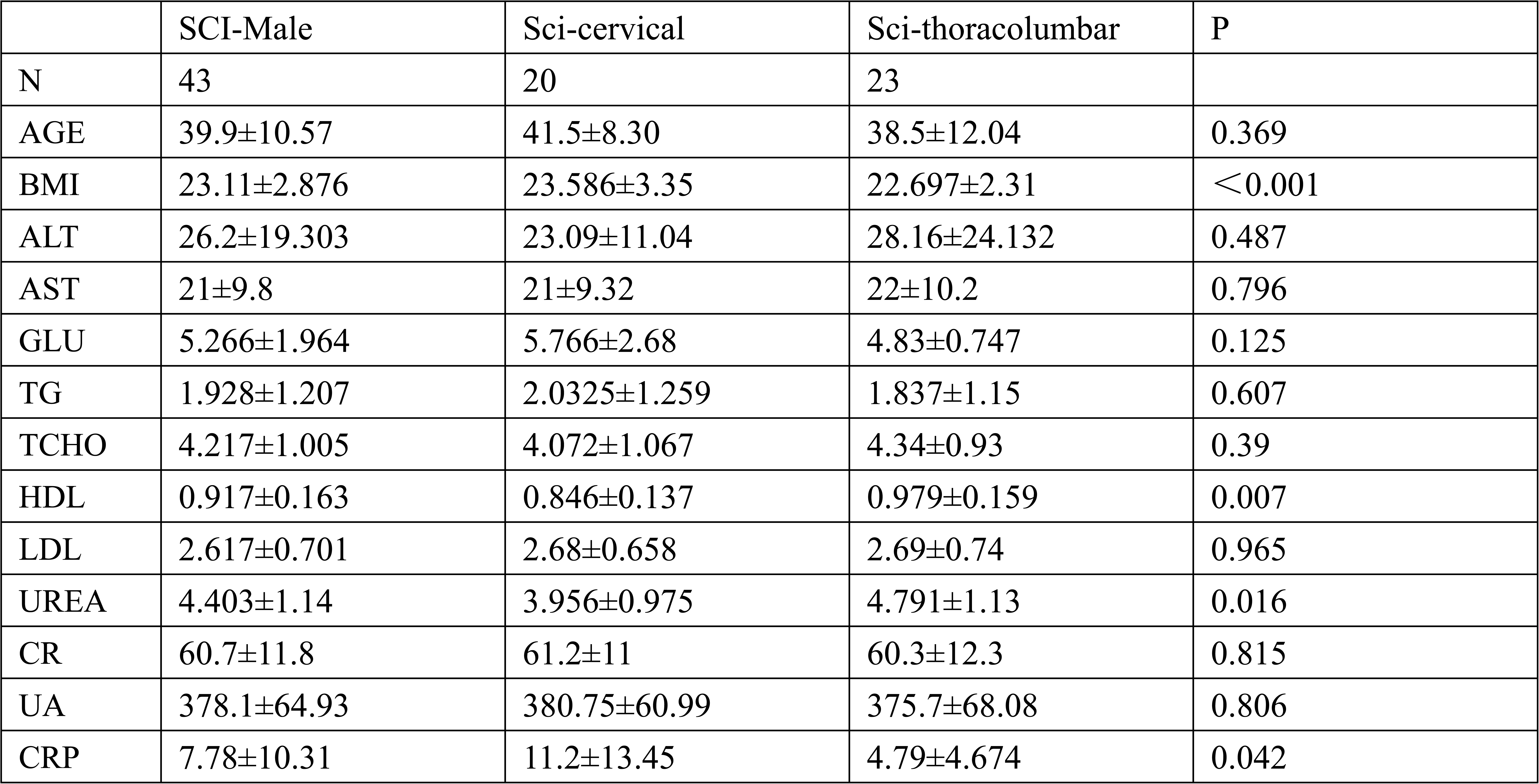
Demographics and serum biomarkers male Patients with chronic traumatic completed SCI

|  | SCI-Male | Sci-cervical | Sci-thoracolumbar | P |
| --- | --- | --- | --- | --- |
| N | 43 | 20 | 23 |  |
| AGE | 39.9±10.57 | 41.5±8.30 | 38.5±12.04 | 0.369 |
| BMI | 23.11±2.876 | 23.586±3.35 | 22.697±2.31 | <0.001 |
| ALT | 26.2±19.303 | 23.09±11.04 | 28.16±24.132 | 0.487 |
| AST | 21±9.8 | 21±9.32 | 22±10.2 | 0.796 |
| GLU | 5.266±1.964 | 5.766±2.68 | 4.83±0.747 | 0.125 |
| TG | 1.928±1.207 | 2.0325±1.259 | 1.837±1.15 | 0.607 |
| TCHO | 4.217±1.005 | 4.072±1.067 | 4.34±0.93 | 0.39 |
| HDL | 0.917±0.163 | 0.846±0.137 | 0.979±0.159 | 0.007 |
| LDL | 2.617±0.701 | 2.68±0.658 | 2.69±0.74 | 0.965 |
| UREA | 4.403±1.14 | 3.956±0.975 | 4.791±1.13 | 0.016 |
| CR | 60.7±11.8 | 61.2±11 | 60.3±12.3 | 0.815 |
| UA | 378.1±64.93 | 380.75±60.99 | 375.7±68.08 | 0.806 |
| CRP | 7.78±10.31 | 11.2±13.45 | 4.79±4.674 | 0.042 |

The rarefaction curves showed clear asymptotes and the Good’s coverage for the observed OTUs was 99.88%, which together indicate a near-complete sampling of community (Fig.5A). No significant difference in OTU abundance (ace, chao index) was observed between the three groups. Significant difference in genus abundance chao index was observed between quadriplegia group and paraplegic group (P=0.02922), healthy male group and paraplegic group (P=0.02919), those indicates a difference in community richness in the two groups (Fig.5B). The Simpson index of health male group showed a significant high than paraplegic group (P=0.04094) in genus level, this indicates a decrease in intestinal flora diversity in patients with paraplegic spinal cord injury (Fig.5C and Supplementary File 5-6).

**Figure 5.**
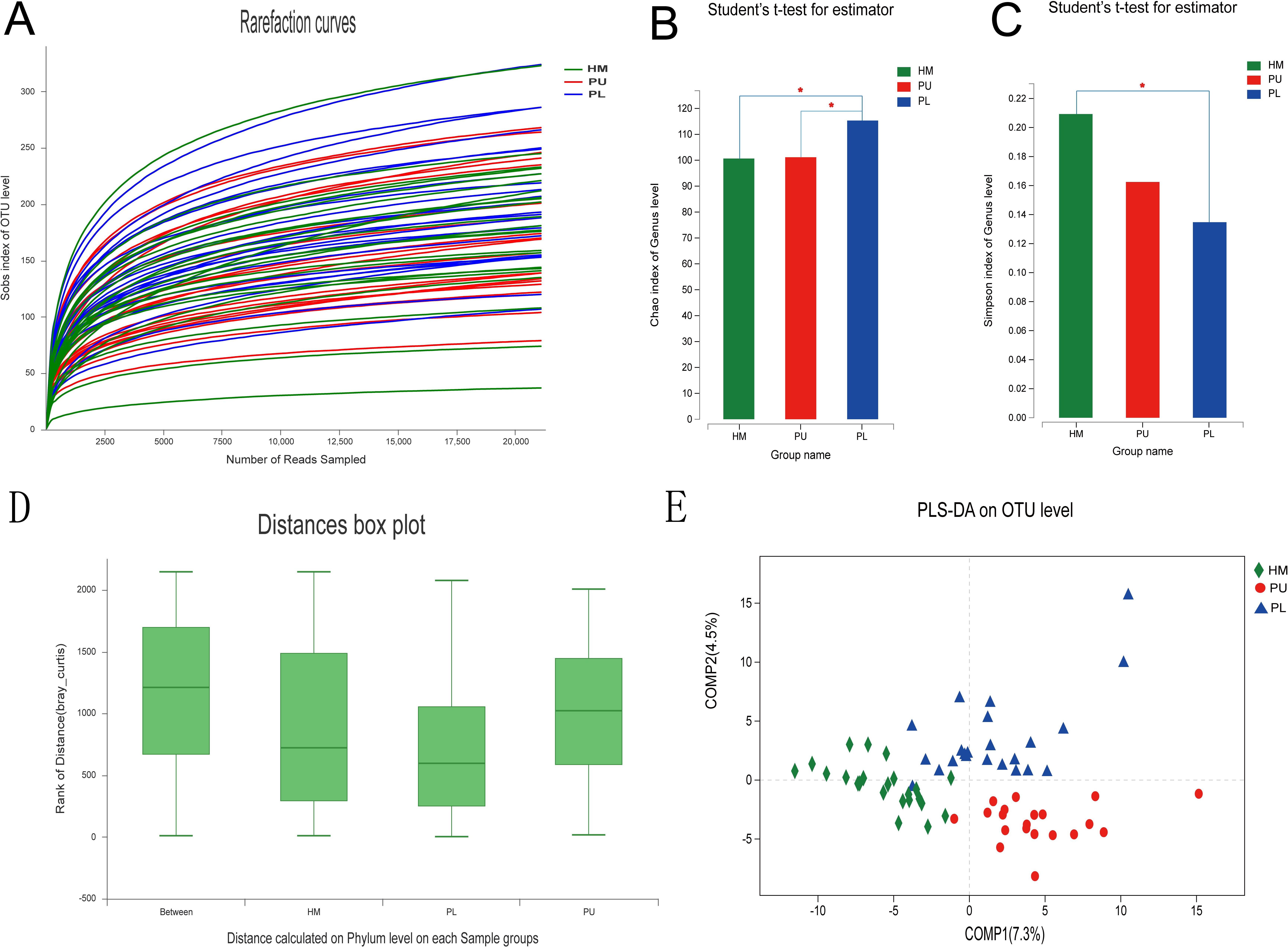
Diversity and taxonomic analysis in the health male, quadriplegia and paraplegic SCI groups. A. Sobs index of Rarefaction curves for healthy male, quadriplegia and paraplegic groups of samples based on OTUs detected using a similarity threshold of 97%. B. Significant difference in genus chao index(B) and Simpson index (C) were observed between the three populations(P< 0.05). D. ANOSIM/Adonis of beta-diversity analysis revealed significant differences in the structure of gut microbiota among the three groups (p=0.001, r^2^=0.233)in phylum level. E.PLS-DA revealed that there were significant differences in bacterial community composition between three groups on OTUs, phylum and genus level.

ANOSIM/Adonis of beta-diversity analysis revealed significant differences in the structure of gut microbiota among the three groups (p=0.001, r^2^=0.233) in phylum level (Fig.5D and Supplementary File 7). PLS-DA revealed that there were significant differences in bacterial community composition between three groups on OTUs, phylum and genus level (Fig.5E).

STAMP analysis indicates there were 8 OTUs showed a significant difference (P<0.05) among three groups (Welch’s t-test)in top 15 OTUs. There were 8 of top 15 genus showed a significant difference (P<0.05) among three groups (Welch’s t-test). The abundance of Firmicutes in paraplegic group and healthy male group were significant high than the quadriplegia group (P= 0.0251 和 P= 0.0185,One-way ANOVA test). In top 15 genus, the abundance of Bacteroides, Faecalibacterium, Blautia, Prevotella_9, Phascolarctobacterium, Parabacteroides, (Eubacterium)_rectale showed a significant difference between the three groups (P< 0.05, One-way ANOVA test)(Fig.6).

**Figure 6.**
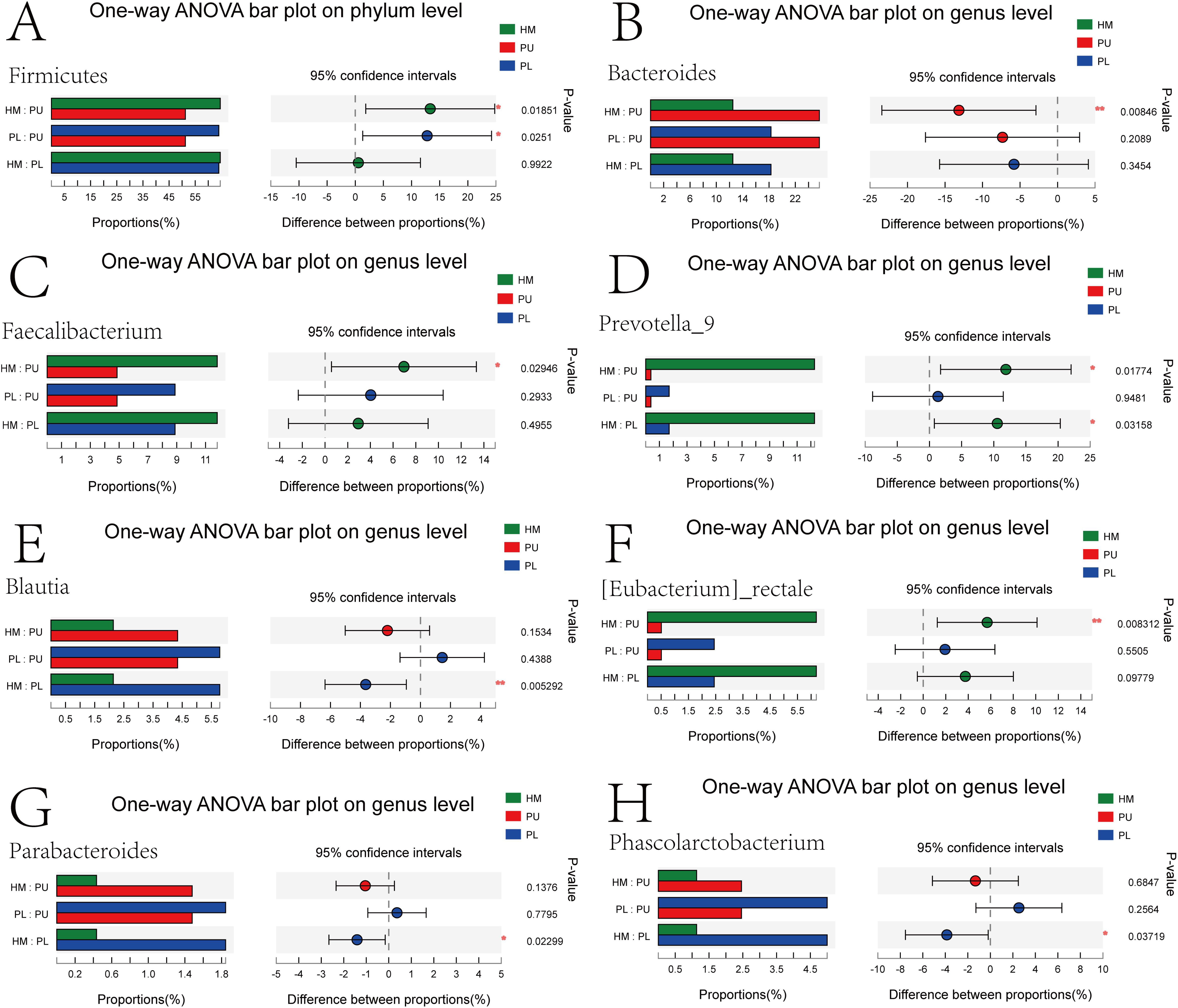
STAMP analysis indicates the significant difference phylum(A) and genus (B-H) among three groups.

The selected 16 environmental factors were: BMI, ALT, AST, GLU, TG, TCHO, HDL, LDL, UREA, CR, UA, AGE, COURSE, CRP, NBD-score, Defecation time for RDA analysis in quadriplegia group and paraplegic group. One-way ANOVA showed statistically significant differences in BMI, HDL, UREA, APOA1and defecation time between the two groups (Table3).

RDA/CCA showed that GLU (p=0.014, r^2^=0.1969) HDL (p=0.009, r^2^=0.2274) CR (p=0.006, r^2^=0.2306), significantly affected bacterial composition in phylum level; In OTU level, TG (p=0.042, r^2^=0.2192), CR (p=0.007, r^2^=0.2388), Defecation time (p=0.022, r^2^=0.2009) significantly affected bacterial composition; In genus level, HDL (p=0.001, r^2^=0.4675), CR (p=0.001, r^2^=0.3209) significantly affected bacterial composition.

Correlation heatmap analysis of different environmental factors on the community composition of quadriplegia and paraplegic groups showed that Alistipes were negatively correlated with defecation time (Pearson r=-0.363,p=0.017), negatively correlated with course (Pearson r=-0.375, p=0.013). In phylum level: Firmicutes was negatively correlated with CRP (Pearson r=-0.491, p=0.001), positively correlated with HDL (Pearson r=0.419, p=0.005). At the genus and OUT levels, the results indicated that the HDL, LDL, CR, UA, AGE have an effect on the intestinal microbiota of two groups (Fig7, Supplementary File 8-9).

**Figure 7.**
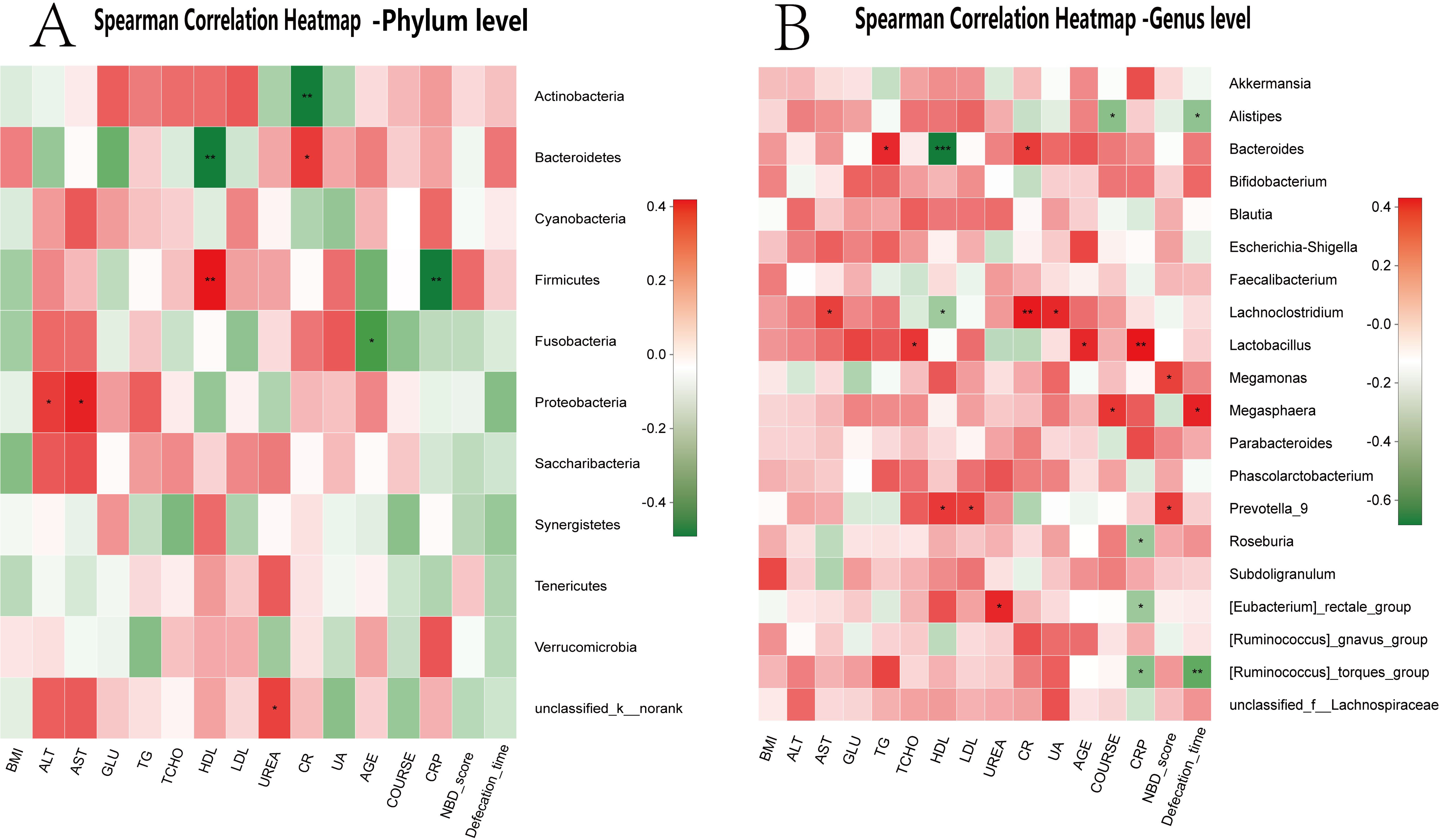
Correlation heatmap analysis of different environmental factors on the community composition of the quadriplegia and paraplegic groups in phylum level (A) and genus level(B).

## Discussion

In this study, after verified the differences in gut microbiota between healthy adult males and females, we compared the gut microbiome between healthy adult males and male patients with chronic traumatic complete SCI. The neurogenic bowel management of SCI patients in our center were firstly reported through cross-sectional interviews. We try to explore the association between gut microbiota and environmental factors in quadriplegia and paraplegic groups; analyse the correlation between gut microbiota and neurogenic bowel symptoms. The results of neurogenic bowel symptoms in SCI patients were related to some gut microbiota and it may help explain the potential link between gut dysbiosis and NBD symptoms in SCI patients.

Acute traumatic SCI (ASCI) used to appear mostly in adults (21—69 years), the causes of injury were fall from height (37.5%), traffic accidents (26.9%) (32,33). This was also consistent with traffic accidents (37.2%), fall from height (20.9%) in this study. The average age of SCI patients in this study was 39.9 years old, which was in the prime of life and was more susceptible to accidental injuries such as high-energy trauma. After chronic course, the most common complication those patients hope to resolve was NBD. Gut microbiota may be a potential method to improve this problem.

Li et al has reported the male/female ratio of ASCI was 3.1/1(32), most of the patients with chronic traumatic complete SCI admitted to our center were males, so we chose male patients as our research objects. Elin O et al had reported the different in gut microbiota between male and female before (22-23). A recent study by Haro et al highlighted differences between men and women in the luminal microbial population. Their results suggest that these differences may be influenced by the grade of obesity (34). The abundance of Bacteroides was higher in females than in males was same with Haro’s research (34). In fact, gut microbiota composition seems to be more influenced by ambient and dietary cues than by genetic factors and inter-individual heterogeneity (35,36). We compared the gut microbiota of healthy males and male SCI patients to eliminate the gender impact.

Julia et al illustrated the practice and outcomes of bowel care in the community of individuals with SCI in Malaysia (37). R Yasmeen had reported that 43 of 50 adult patients with SCI in Pakistan gave a history of occasional or regular fecal incontinence (38). The prevalence of NBD in SCI patients was 80% in previous study, 97.3% of motor complete SCI patients had chronic NBD complaints in their study (10), which was similar with patients had a constipation in our study. The patients in our study had to spend much time on defecation in their long course of SCI to deal with NBD.

Injury level has been shown not to be related to gastro-intestinal complaints in SCI patients detected no significant relation between gastro-intestinal symptom prevalence and SCI level in their study (39,40,41). Most patients with quadriplegia have no active exercise capacity but paraplegic patients can complete all upper limb movements. The autonomic nervous system that supports the gastrointestinal tract in quadriplegia SCI patients remains relatively intact and has less effect on the function of the gastrointestinal tract. Patients with paraplegia had damaged the sympathetic center or defecation center would had a greater impact on intestinal function. This may explain the different between the two groups.

Surveys among the SCI population often rank colorectal, bladder and sexual dysfunction as significant obstacles and prioritize recovery of bowel function above the ability to walk (42,43,44). In this study, the most common top 3 complications that patients wanted to solved were NBD, neurogenic bladder, sexual dysfunction, the gut microbiota may be a potential solution strategy.

Rajilić-Stojanović observed an increase in the phylum Bacteroidetes, which was reflected by increased Bacteroides at the genus level. The phylum Bacteroidetes encompasses a diverse and abundant group of gram-negative commensal bacteria in the gut (45). The major outer membrane component of gram-negative bacteria is lipopolysaccharide (LPS), which is capable of triggering systemic inflammation and the release of pro-inflammatory cytokines after translocation from the gut to systemic circulation (46). In our study, we found the Bacteroides was significant high in SCI group and negatively correlated with HDL, especially enriched in quadriplegia group. HDL levels are positively correlated with amount of exercise, and lack of exercise can result in lower HDL levels (47). Reduced exercise in quadriplegia patients made the lowers HDL levels, and the Bacteroides were increased in quadriplegia patients. Those indicated that Bacteroides may be a harmful flora and was associated with the exercise v and serum HDL levels.

Nicholas M. Vogt had reported that Dialister showed the strongest correlations in non-demented participants, with greater abundance of these bacteria associated with less Alzheimer’s disease (AD) pathology, suggesting these bacterial taxa may be protective against development or progression of AD pathology (48). In our study, the Dialister was significant high in health males, and these bacteria were negatively correlated with UA, LDL, TG and TCHO. Those elevated serum markers represent high blood lipids, which are harmful to the health. We found that the decreased Dialister in SCI patients may aggravated the symptoms of the NBD.

Ming et al reported the positive association between bean consumption and the Megamonas genus discovered in their study may implicate Megamonas as a beneficial microbe(49). The abundance of Megamonas was decreased in SCI group and was negatively correct with GLU, positively with NBD scores, implicate that Megamonas may have a positive effect on the body in terms of carbohydrate metabolism. The decreased Megamonas exacerbates NBD symptoms. We found that in bloating group the relative abundance of Megamonas had a significant high than without bloating group, this may be due to the fact that some carbohydrates in food cannot be digested and absorbed by intestinal digestive enzymes, but they can be metabolized by Megamonas in the large intestine, that producing gas and causing bloating, exacerbates NBD symptoms.

Genera of Alistipes were already reported to be altered in irritable bowel syndrome, animal-based diet or vegetables consumption (50,51,52). Alistipes were associated with the phenotype of frequently recurrent abdominal pain (50). We found the relative abundance of Alistipes was significant decreased in bloating group, and Alistipes were negatively correlated with defecation time, this may be a factor that affect the defecation time between quadriplegia and paraplegic group because the abundance of Alistipes and the defecation time in quadriplegia group were significant high than the paraplegic group. But there is still a big gap in knowledge to explain biochemical and functional roles of Alistipes.

Members of the Bifdobacterium genus, are an important bacterial inhabitant of the human gut across the lifespan, and their beneficial health effects have been well-documented (53,54). But certain species of Bifdobacterium are associated with decreased intestinal permeability. In our study the abundance of Bifdobacterium and Bacteroides were increased in SCI group, it may because the increased bacterial translocation effect of Bacteroides played a more important role than Bifdobacterium. The relative abundance of Bifdobacterium was significant high in constipation group in our study, we thought in the chronic course of constipation symptom, the patients may have used the probiotics which including Bifdobacterium. More investigation is needed to determine the interaction of these bacterial.

We also examined the association between serum biomarkers and the gut microbiota. Prevotella is considered a beneficial microbe, but it is also linked with chronic inflammatory conditions (55,56,57,58). Our study found a decreased Prevotella level in SCI group and negatively correct with GLU, which implicate that Prevotella have a positive effect on the body in terms of carbohydrate metabolism supporting Prevotella as a beneficial microbe.

A strong point of this study was the inclusion of only complete SCI male patients. This approach excluded probable confounding effects of gender and residual nerves on gut functions, and therefore other incomplete injuries were not included in this study. Individual diet-associated flora differences could not be determined and remains as a major weakness of this study. The continuing analyses of genomic and metagenomic changes in gut microbiota will allow scientists to map the dynamic patterns of dysbiosis caused by SCI.

Only male SCI patients were enrolled in our study, and future studies should include female patients to identify gender disparities. Further work, including animal experiments and longitudinal human studies, will be needed to determine the cause-effect relationship between gut microbiota and SCI. Determining the role of gut microbiota in the progression or maintenance of SCI may lead to novel interventional approaches that alter or restore healthy gut bacterial composition, or identification of microbial metabolites that are protective against SCI.

In conclusion, this study presents a comprehensive landscape of gut microbiota in adult male patients with traumatic complete SCI and documents their neurogenic bowel management. We found a difference in fecal flora between healthy adult males and females; Dysbiosis of SCI patients was correlation with Serum biomarkers and NBD symptoms.

## Materials and methods

### Ethics statement

Approval of hospital ethics committee was obtained before commencing the study.

### Patients and controls

A total of 43 chronic traumatic complete SCI male patients (20 with quadriplegia and 23 with paraplegia) in our center from March 2017 to October 2017 were enrolled to face-to-face clinical questionnaire survey. Signed informed consent before the assessment, and use “International Spinal Cord Injury Core Data Set”, “International bowel function basic spinal cord injury data set” and “International bowel function extended spinal cord injury data set” to get the NBD symptoms dates (59,60,61).

Patients were included if they met the following criteria: 1) neurologically complete SCI (ASIA grade A) occurring 6 or more months prior to study, 2) 18-60 years of age, 3) traumatic spinal cord injury, 4) male patients. The exclusion criteria: 1) patients who can not cooperate for questionnaire survey, 2) with a history of antibiotic use in the first month before enrollment, 3) patients with diabetes, gastrointestinal system diseases, multiple sclerosis, and immune metabolic diseases.

A total of 43 SCI patients and 48 healthy adults (23 males and 25 females) were enrolled in to collect clinical dates of the subjects and fresh stool specimens, extract fecal genomic DNA, amplify the V3-V4 region of 16S rDNA, and sequence 11mmola MiSeq platform to analyze the gut microbiota of healthy male with female and healthy male with SCI patients.

The healthy control group included criteria: 1) 18-60 years of age, 2) without a history of antibiotics or probiotics use 1 month prior to study, 3) without the history of diabetes, gastrointestinal system diseases, multiple sclerosis, and immune metabolic diseases. All subjects selected before sampling and signing informed consent fully understand the sampling process and research options. To exclude probable effects of diet on microbiota, all patients and healthy subjects were fed with standard hospital food 2 weeks before stool collection.

### 16S Diversity materials and methods

#### Microbial diversity analysis

##### 1. Stool sampling

91 fresh specimens were collected, including 23 healthy male, 25 healthy female, 43 SCI patients. Fresh fecal samples were collected and transferred to the laboratory. Their 200 mg sample was placed in a new 2-mL sterile centrifuge tube, quickly placed on ice, and transferred to a refrigerator-80 °C cryostat for cryopreservation. The entire sampling process is completed in 30 minutes.

##### 2. DNA extraction and PCR amplification

Microbial DNA was extracted from stool samples using the E.Z.N.A.® Stool DNA Kit (Omega Bio-tek, Norcross, GA, U.S.) according to manufacturer’s protocols. The V3-V4 region of the bacteria 16S rRNA gene were amplified by PCR (95 °C for 2 min, followed by 25 cycles at 95 °C for 30 s, 55 °C for 30 s, and 72 °C for 30 s and a final extension at 72 °C for 5 min) using primers 338F 5’-ACTCCTACGGGAGGCAGCA-3’ and 806R 5’- GGACTACHVGGGTWTCTAAT-3’. PCR reactions were performed in triplicate 20 μL mixture containing 4 μL of 5 × FastPfu Buffer, 2 μL of 2.5 mM dNTPs, 0.8 μL of each primer (5 μM), 0.4 μL of FastPfu Polymerase, and 10 ng of template DNA.

##### 3. Illumina MiSeq sequencing

Amplicons were extracted from 2% agarose gels, purified by using the AxyPrep DNA Gel Extraction Kit (Axygen Biosciences, Union City, CA, U.S.), and quantified by using QuantiFluor™ -ST (Promega, U.S.). Purified amplicons were pooled in equimolar and paired-end sequenced (2 × 300 bp) on an Illumina MiSeq platform according to the standard protocols. The raw reads were deposited into the NCBI Sequence Read Archive database (Accession Number: SRP158549).

##### 4. Processing of sequencing data

Raw fastq files were quality-filtered by Trimmomatic and merged by FLASH with the following criteria: (i) The reads were truncated at any site receiving an average quality score < 20 over a 50 bp sliding window. (ii) Sequences whose overlap being longer than 10 bp were merged according to their overlap with mismatch no more than 2 bp. (iii) Sequences of each sample were separated according to barcodes (exactly matching) and Primers (allowing 2 nucleotide mismatching) and reads containing ambiguous bases were removed.

Operational taxonomic units (OTUs) were clustered with 97% similarity cutoff using UPARSE (version 7.1) and chimeric sequences were identified and removed using UCHIME. The taxonomy of each 16S rRNA gene sequence was analyzed by RDP Classifier algorithm against the Silva (SSU123) 16S rRNA database using confidence threshold of 70% Roche 454 (Roche, Switzerland) high-throughput sequencing of the PCR products was performed by Shanghai Majorbio Biological Technology Co. Ltd., Shanghai, China.

### Bioinformatic and statistical analysis

Sequencing reads were processed using QIIME (version 1.9.0), and included additional quality trimming, demultiplexing, and taxonomic assignments. Profiling of predictive urine microbiota was analyzed by using PiCRUSt based on 13 August 2013 Greengenes database (62). KW rank sum test and pairwise Wilcoxon test were used for the identification of the different markers, and LDA was used to score each feature in the LEfSe analysis. Index of alpha diversity was calculated with QIIME based on sequence similarity at 97%. Beta diversity was measured by unweighted UniFrac distance, which was also calculated by QIIME. Hierarchical clustering was performed, and a heatmap was generated using a Spearman’s rank correlation coeffcient as a distance measure and a customized script developed in the R statistical package. The output file was further analyzed using Statistical Analysis of Metagenomic Profiles software package (version 2.1.3) (63).

Statistical analysis was performed using the SPSS data analysis program (version 21.0) and Statistical Analysis of Metagenomic Profiles software. For continuous variables, independent t-test, Welch’s t-test, White’s nonparametric t-test, and Mann-Whitney U-test were applied. For categorical variables between groups, using either the Pearson chi-square or Fisher’s exact test, depending on assumption validity. For taxon among subgroups, ANOVA test was applied (Tukey-Kramer was used in Post-hoc test, Effect size was Eta-squared) with Benjamini-Hochberg FDP false discovery rate correction (63,64). All tests of significance were two-sided and p <0.05.

## Abbreviations

SCI: spinal cord injury
NBD: neurogenic bowel dysfunction
GLU: glucose
HDL: high density lipoprotein
LDL: low density lipoprotein
UA: Uric acid
CR: Creatinine
CPR: C-reactive protein
OUTs: Operational taxonomic units
PLS-DA: Partial least squares discrimination analysis
BMI: Body mass index
APOA1: Apolipoprotein A1
APOB: Apolipoprotein B
ALT: Alanine transaminase
AST: Aspartate transaminase
TG: Triglyceride
TCHO: Total Cholesterol
LPA: lipoprotein A
NEFA: non-esterified fatty acid
HCY: homocysteine
ASCI: acute spinal cord injury
LPS: lipopolysaccharide
AD: Alzheimer’s disease
HM: healthy male
FM: healthy female
PU: quadriplegia SCI patient
PL: paraplegic SCI patient.

## Acknowledgements

This work was supported by the Special Fund for Basic Scientific Research of Central Public Research Institutes, grant number: 2016cz-1, 2018cz-8, and Beijing Municipal Science and Technology Commission (BSTC, No. Z171100001017076).

## Supplementary File Figure legends

Supplementary File 1:

Sobs index of Rarefaction curves for healthy male and female groups of samples based on OTUs detected using a similarity threshold of 97%.

Supplementary File 2:

ANOSIM/Adonis on OUT, phylum and genes level revealed significant differences in the structure of gut microbiota among the healthy male and SCI groups (p<0.05).

Supplementary File 3

Correlation heatmap analysis chart of different environmental factors on the community composition of healthy male and SCI groups in genus level.

Supplementary File 4

Alpha-diversity index inter-group difference test chart between healthy male, quadriplegia and paraplegic SCI cohorts in OTUs level.

Supplementary File 5:

Alpha-diversity index inter-group difference test chart between quadriplegia and paraplegic SCI cohorts in genus level.

Supplementary File 6:

Alpha-diversity index inter-group difference test chart between healthy male and paraplegic SCI cohorts.

Supplementary File 7:

ANOSIM/Adonis distances box plot on phylum level revealed significant differences in the structure of gut microbiota among the healthy male, quadriplegia and paraplegic SCI groups (p< 0.05).

Supplementary File 8

Correlation heatmap analysis chart of different environmental factors on the community composition of quadriplegia and paraplegic SCI groups in phylum level.

Supplementary File 9

Correlation heatmap analysis chart of different environmental factors on the community composition of quadriplegia and paraplegic SCI groups in genus level.

